# A unique predator in a unique ecosystem: modelling the apex predator from the late cretaceous crocodyliform-dominated fauna in brazil

**DOI:** 10.1101/843334

**Authors:** Felipe C. Montefeltro, Stephan Lautenschlager, Pedro L. Godoy, Gabriel S. Ferreira, Richard J. Butler

## Abstract

Theropod dinosaurs were relatively scarce in the Late Cretaceous ecosystems of southeast Brazil. Instead, hypercarnivorous crocodyliforms known as baurusuchids were abundant and probably occupied the ecological role of apex predators. Baurusuchids exhibited a series of morphological adaptations hypothesised to be associated with this ecological role, but quantitative biomechanical analyses of their morphology have so far been lacking. Here, we employ a biomechanical modelling approach, applying finite element analysis (FEA) to models of the skull and mandibles of a baurusuchid specimen. This allowed us to characterise the craniomandibular apparatus of baurusuchids, as well as to compare the functional morphology of the group to that of other archosaurian carnivores, such as theropods and crocodylians. Our results support the ecological role of baurusuchids as specialised apex predators in the continental Late Cretaceous ecosystems of South America. With a relatively weak bite force (∼600 N), baurusuchids’ predation strategies likely relied on other morphological specializations, such as ziphodont dentition and strong cervical musculature. Comparative assessments of the stress distribution and magnitude of scaled models of other predators (the theropod *Allosaurus fragilis* and the living crocodylian *Alligator mississippiensis*) consistently show different responses to loadings under the same functional scenarios, suggesting distinct predatory behaviours for these animals. The unique selective pressures in the arid to semi-arid Late Cretaceous ecosystems of southeast Brazil, which were dominated by crocodyliforms, possibly drove the emergence and evolution of the biomechanical features seen in baurusuchids, which are distinct from those previously reported for other predatory taxa.

## INTRODUCTION

In nearly all known continental Cretaceous ecosystems worldwide, the dominant hypercarnivores and apex predators were theropod dinosaurs (Lloyd *et al*. 2008; Benson *et al*. 2013; Zanno & Mackovicky 2013). However, in the Late Cretaceous ecosystems of Brazil, theropods were exceptionally scarce. Instead, the putative dominant apex predators were a group of large, terrestrial crocodyliforms, the baurusuchids (Riff & Kellner 2011; Godoy *et al*. 2014). Baurusuchids are phylogenetically included within Notosuchia, a group of highly diverse crocodyliforms which thrived mainly in Gondwana during the Cretaceous (Pol & Leardi 2015; Mannion *et al*. 2015). Exhibiting a wide range of morphological variation, from gracile omnivores to pug-nosed herbivores, notosuchians contributed significantly to the highest peak of morphological disparity experienced by crocodyliforms across their evolutionary history (Wilberg 2017; Godoy 2019; Godoy *et al*. 2019; Melstrom & Irmis 2019).

Although present in other parts of Gondwana, most baurusuchid species (ca. 80%) are found in the Late Cretaceous rocks of the Bauru Group, in southeast Brazil (Carvalho *et al*. 2005; Godoy *et al*. 2014; Montefeltro *et al*. 2011). The Bauru Group palaeoecosystem witnessed an extraordinary abundance of notosuchians, with nearly 30 species described so far. Dinosaurs were also present, but their fossil record in this rock sequence is relatively poor (Montefeltro *et al*. 2011; Godoy *et al*. 2014). Within this crocodyliform-dominated ecosystem, baurusuchids formed the likely apex predators. Baurusuchids exhibited a series of morphological adaptations hypothesised to be associated with their role as hypercarnivores, possibly achieved via heterochronic transformations, such as hypertrophied canines, a reduced number of teeth, and dorsoventrally high skulls (Montefeltro *et al*. 2011; Riff & Kellner 2011; Godoy *et al*. 2018). However, quantitative assessments of the palaeobiology of baurusuchids are lacking, and the data supporting their role as apex predators is primarily derived from broad generalizations and the faunal composition of the Bauru palaeoecosystem (Riff & Kellner 2011; Godoy *et al*. 2014).

Here, we employ a biomechanical modelling approach to test the hypothesis that the functional morphology of their skulls allowed baurusuchids to outcompete other contemporary archosaurian carnivores. Using finite element analysis (FEA), we characterize the baurusuchid skull biomechanically and quantify functional similarities and differences between baurusuchids, theropod dinosaurs and living crocodylians. We also calculate bite forces, simulate functional scenarios and conduct bending tests to reveal biomechanical properties of the baurusuchid skull and to understand how this group dominated the unique ecosystems present during the Cretaceous in Brazil.

## MATERIALS AND METHODS

### Specimens

The baurusuchid specimen modelled for the present study is a complete skull with lower jaws, referred to *Baurusuchus pachecoi* (LPRP/USP 0697 Laboratório de Paleontologia USP-RP) and collected in Jales, Brazil (Adamantina Formation, Bauru Group; Montefeltro 2019). For comparison, we modelled a specimen of the theropod dinosaur *Allosaurus fragilis* (MOR 693, Museum of the Rockies, Bozeman) and one specimen of *Alligator mississippiensis* (OUVC 9761, Ohio University Vertebrate Collections) (see Rayfield *et al*. 2001, Witmer & Ridgely 2008 for scanning details). *Allosaurus fragilis* was chosen based on its medium size when compared to other theropods, which is equivalent to the putative size of the theropods from the Adamantina Formation, for which no complete craniomandibular material is currently known. Furthermore, *Allosaurus* has been proposed to be functionally similar to abelisaurids, the most commonly found theropods in the Bauru Group (Sakamoto 2010). The choice of *Alligator mississippiensis* (as a living representative of the crocodyliform lineage) was made because this is a model organism for herpetological and functional studies (Guillette *et al*. 2007; Farmer & Sanders 2010; Reed *et al*. 2011). For the subsequent FEA, existing 3D models of *Allosaurus fragilis* and *Alligator mississippiensis* from previous studies were used (Rayfield *et al*. 2001; Witmer & Ridgely 2008; Lautenschlager 2015). The *Baurusuchus pachecoi* skull was scanned in a Toshiba Aquilion Prime machine, at “Hospital das Clínicas de Ribeirão Preto”, Brazil. The scan resulted in 1917 projections, generating 1.187 slices (thickness of 0,5 cm), voltage of 120 kV, and current of 150 μA. The segmentation of bones was achieved with Amira 5.3.

### FEA

The 3D models of all specimens, including skulls and mandibles, were imported into Hypermesh 11 (Altair Engineering) for the generation of solid tetrahedral meshes (consisting of approximately 1,000,000 elements per model). For the *Alligator* and the baurusuchid models, material properties for bone and teeth were assigned based on values for *Alligator mississippiensis* (bone: E = 15.0 GPa, ν = 0.29, teeth: E = 60.4 GPa, ν = 0.31; Porro *et al*. 2011; Sellers *et al*. 2017), whereas for the *Allosaurus* model, values were derived from studies on theropods (bone: E = 20.0 GPa, ν = 0.38, teeth: E = 60.4 GPa, ν = 0.31; Rayfield *et al*. 2001, 2011). To exclude the possibility of different results due to distinct material properties we also conducted an FEA on the *Allosaurus* model using the same bone and teeth properties assigned to the crocodyliform models. All material properties in the models were assigned in Hypermesh and treated as isotropic and homogeneous.

Intrinsic scenarios for the baurusuchid, *Allosaurus fragilis* and *Alligator mississippiensis*, were simulated for the skull and lower jaw models, using a simplified jaw adductor muscle-driven biting. The adductor muscle forces of the baurusuchid were estimated using the attachment area for each muscle, based on previous works on extant and extinct crocodyliforms (Holliday & Witmer 2009; Holliday *et al*. 2013). The adductor chamber reconstruction of the dinosaur and crocodylian was based on previously published data for the muscle arrangements for both taxa (Rayfield *et al*. 2001, 2011; Porro *et al*. 2011; Sellers *et al*. 2017). The attachment areas measured for the three taxa were used as a proxy for physiological cross-section area, which was then multiplied by an isometric muscle stress value of 25.0 N/cm2 (Porro *et al*. 2011). Although this isometric muscle stress is on the lower margin of the range of values reported for vertebrate muscles (e.g. 32N/cm2 and 35N/cm2) it was selected here due to the relatively close phylogenetic position of baurusuchids to modern crocodilians. However, the calculated bite force would be only slightly (10-15%) higher using different values for isometric muscle stress. Three intrinsic scenarios were analysed to estimate the muscle-driven biting force in the baurusuchid, the bilateral bite scenario for the skull and lower jaw, maxillary and dentary unilateral bite scenario skull and lower jaw, and premaxillary unilateral bite scenario. One intrinsic scenario was analysed for both *Allosaurus fragilis* and *Alligator mississippiensis*: the maxillary and dentary unilateral bite scenarios. For each intrinsic scenario in all taxa, constraints were placed on nodes at the craniomandibular articular surfaces. Each node was constrained in all directions (x, y, z). For the skulls, three nodes were constrained on the occipital condyle, and two nodes on each quadrate articular surface. For the lower jaws, three (baurusuchid) or four (*Allosaurus* and *Alligator*) nodes on each glenoid were constrained. To estimate the biting force of the baurusuchid, nodes were constrained at the tip of the teeth to measure the reaction force caused by the modelled adductor muscles and the same approach was used for the other two taxa. In unilateral scenarios, the tip of one tooth was constrained, while in bilateral scenarios the tip of the teeth on both sides were constrained. For the baurusuchid, the constrained teeth were PM3, M2 and D4; for *Allosaurus fragilis*, M3 and D5; for *Alligator mississippiensis*, M4 and D4.

To investigate the craniomandibular biomechanical properties in alternative load assignments, bending scenarios were also tested for the baurusuchid skull and mandible models: unilateral bending, bilateral bending, pull-back, head-shake and head-twist. The loading applied for each scenario was based on the approximation of the greatest bite force obtained from the intrinsic scenario (600 N; see results below). All loadings in the unilateral bending scenario were applied to one node, perpendicular to the occlusal planes on one of the following teeth: D1, D4, D9, PM2, PM3, M2 and M4. Bilateral bending scenarios were tested with the same conditions as the unilateral ones, but with two vectors of 300 N applied to each canine at the M4 and the D4. The head-shake scenario was tested with two vectors of 300 N pointing to the same direction, one on one node on the labial surface of left M2/D4 and the other on one node on the lingual surface of right M2/D4. For the pull-back, the load force of 600 N was applied to one node at crown midheight over the distal carina of the caniniform teeth (D4, PM3 and M2). For the head twist, the loadings were applied to two opposite vectors of 300 N in each model. One loading vector was applied to one node at the tip of the maxillary (M2) or dentary (D4) caniniform tooth, and another loading vector on the opposite side on the dorsal surface of the maxilla, or ventral surface of the dentary respectively.

Four bending scenarios were also tested in the skull and lower jaws of *Allosaurus fragilis* and *Alligator mississippiensis*, for comparison. Unilateral and bilateral bending were simulated to the comparable positions of the tested in the baurusuchid. Unilateral bending was tested in PM2, M3, M16, D1, D4 and D13 for *Allosaurus fragilis*, and PM2, M4, M15, D2, D4 and D15 for *Alligator mississippiensis*. Bilateral bending was also tested in M3 and D5 pairs for the theropod, and M4 and D4 pairs for the crocodylian. For meaningful comparisons of form and function independent of size (Dumont *et al*., 2009), all models used in the bending tests were scaled to the total surface of the baurusuchid specimen. For the bending scenarios, constraints were placed on the same nodes as in the intrinsic scenarios.

## RESULTS

During the bilateral bite scenario, the bite force estimate for the baurusuchid specimen was 252 N for the skull and 578 N for the lower jaw. For the premaxillary unilateral bite scenario, bite force was estimated as 199 N, whereas for both maxillary and lower jaw unilateral bite scenarios, it was 450 N. The distribution and magnitude of the Von Mises stress showed little difference in the intrinsic scenarios for the skull and lower jaw of the baurusuchid (Figure 1). Most of the elements in the skull remained relatively stress-free in the three intrinsic scenarios simulated (average Von Mises stress of 0.46 MPa during the bilateral maxillary biting, 0.50 MPa during the unilateral maxillary biting, and 0.52 MPa during the premaxillary unilateral biting). The quadrate body, the body of the ectopterygoid, and the posterior margin of the pterygoid are the main regions in which stress is present during those simulated scenarios (Figure 1). In the intrinsic scenario for the premaxillary canine bite, there is also increased stress at the anterior margin of the notch between the premaxilla and maxilla, which also extends medially surrounding the notch at the secondary bony palate. As expected, the lower jaws experienced more Von Mises stress than the skull model (average Von Mises stress of 1.93 MPa in the bilateral biting, and 2.01 MPa in the unilateral biting). In both scenarios, the symphyseal region surrounding the canine teeth, and the retroarticular process remained relatively stress-free, and the greatest Von Mises stress is observed on the dorsal surface of the surangular and ventral surface of the angular.

**Figure 1.**
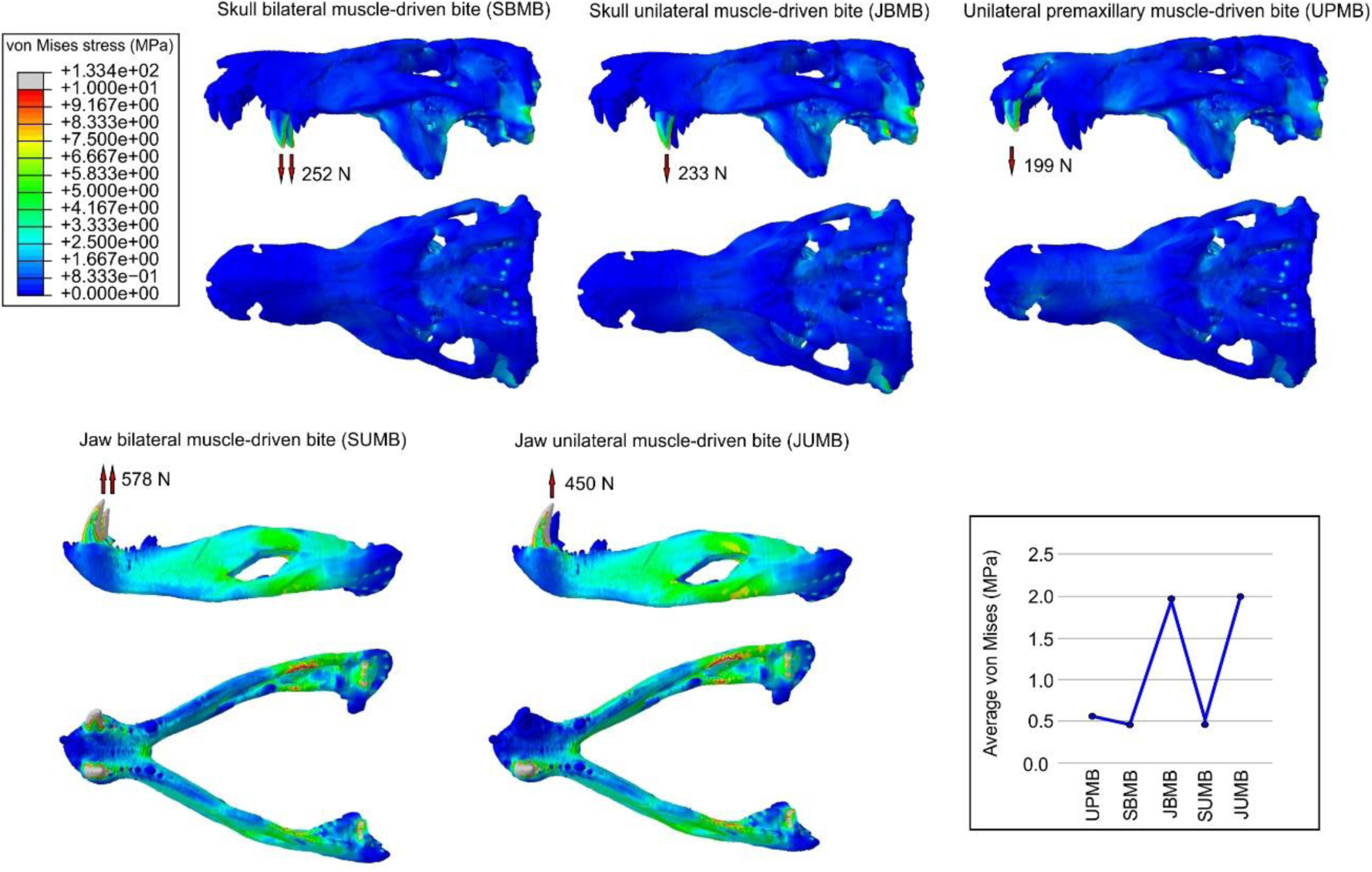
Von Mises stress contour plots from finite elements analysis (FEA) of the baurusuchid specimen (LPRP/USP 0697) for the intrinsic scenarios. Arrows indicate the location of muscle-driven bite forces on models during each scenario, with respective estimated bite force values. Average Von Mises values per scenario are displayed on the bottom right.

Considerable differences were found between the Von Mises stress magnitudes of the skull and lower jaws of the baurusuchid among the different bending scenarios tested (e.g. average values of 0.4 MPa in the skull head twist and of 24.7 MPa in the bilateral biting of the lower jaws). Although variable in magnitude, a general pattern is discernible in the stress distribution in the skull and lower jaws of the baurusuchid (Figure 2). The greatest Von Mises stresses in the skull models are mostly present in the posterior and median portions of the skull, with stress hotspots located on the ventral and lateral regions of the quadrate body, ventral region of the infratemporal bar, and preorbital region (anterior jugal, posterior maxillae, lacrimals, nasal, prefrontals, and anterior frontal). In addition, the areas of maximum Von Mises stress in the premaxillae and maxillae are isolated from each other. This means that when loading is applied to the premaxillary teeth, the maxillae remain relatively stress-free, whereas the dorsal rostrum (premaxilla and nasals) is more stressed. When loading is applied to the maxillary teeth, the premaxillae remain unstressed, and stress is concentrated on the posterior portion of the skull (Figure 2).

**Figure 2.**
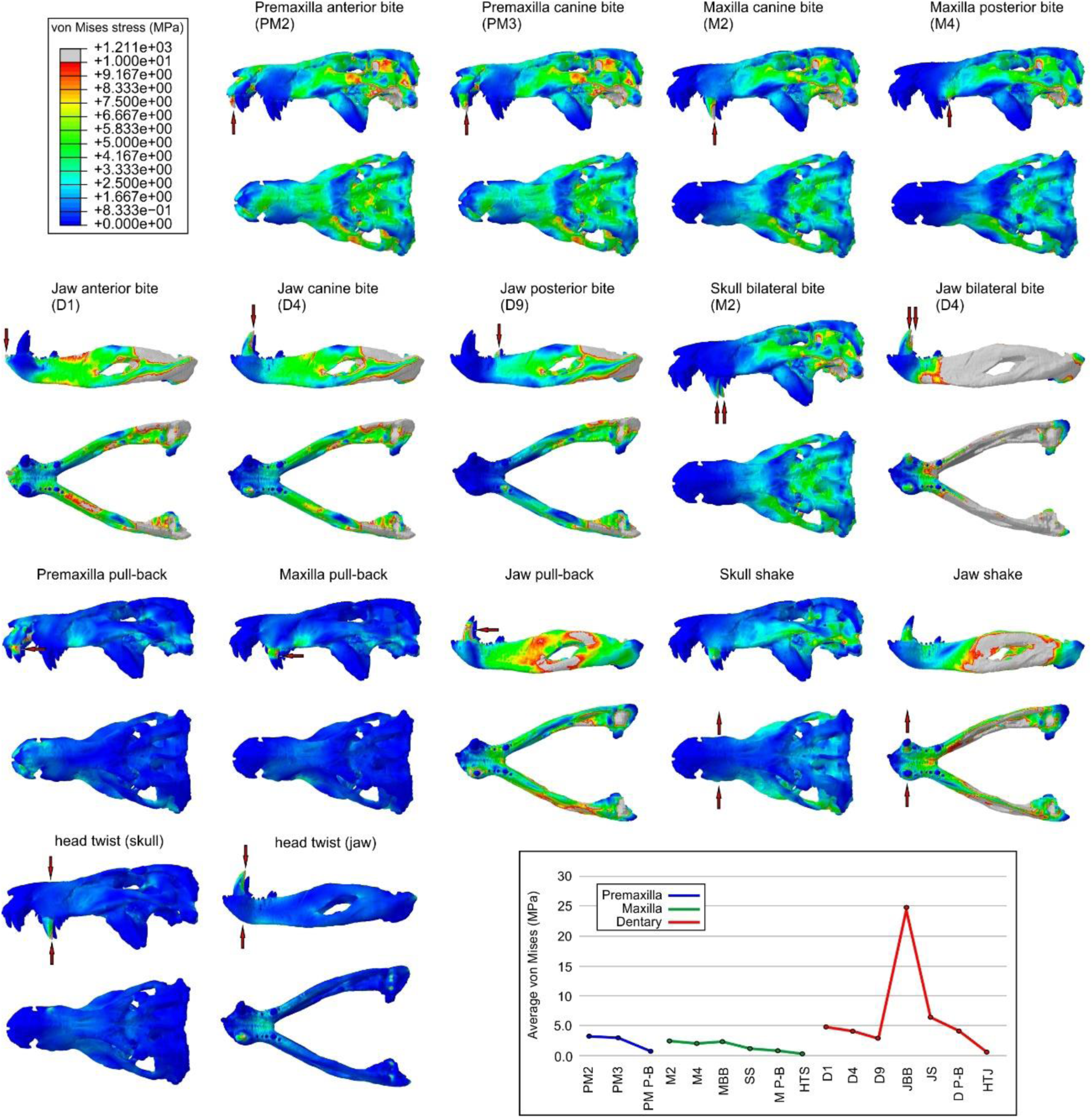
Von Mises stress contour plots from FEA of the baurusuchid specimen LPRP/USP 0697, comparing the stress distribution of skull and mandible models under distinct functional bending scenarios. Arrows indicate the location on the models of the loading vectors for each scenario. Average Von Mises values per scenario are displayed on the bottom right.

The lower jaws also experienced more Von Mises stress than the skull model during the bending tests, and the stress hotspots are more homogeneously distributed, located on the dorsal surface of the surangular, angular and retroarticular process. Two exceptions are the jaw pull back scenario, in which the stress hotspots are located around the mandibular fenestra; and the bilateral bending scenario, in which most of the lower jaw is highly stressed, and only the symphyseal region remains less stressed.

The areas around the maxillary and dentary canines remain relatively stress-free, even during scenarios in which the loadings were applied to the canines (both in the intrinsic scenarios and the bending tests). This is particularly evident for the dentary canine, for which the surrounding bone remains unstressed in all scenarios, including the least optimal scenario of the bilateral bending (Figure 2).

In general, the patterns of Von Mises stress distribution obtained for *Allosaurus* and *Alligator* (Figure 3 and Figure 4) were consistent with previous studies (Rayfield *et al*. 2001; Porro *et al*. 2011). Even considering that the bone properties assigned to the *Allosaurus* are slightly different from the other models, it did not substantially change the results obtained from this taxon. Considering the intrinsic scenarios, the measured average Von Mises stress is similar during maxillary unilateral biting (average Von Mises stress of 0.72 MPa for *Allosaurus* and 0.62 MPa for *Alligator*). The pattern of stress distribution observed in the models of the *Alligator* are much closer to the observed in the baurusuchid than to the *Allosaurus*, perhaps related to the phylogenetic proximity reflected in the cranial architecture of both crocodyliforms.

**Figure 3.**
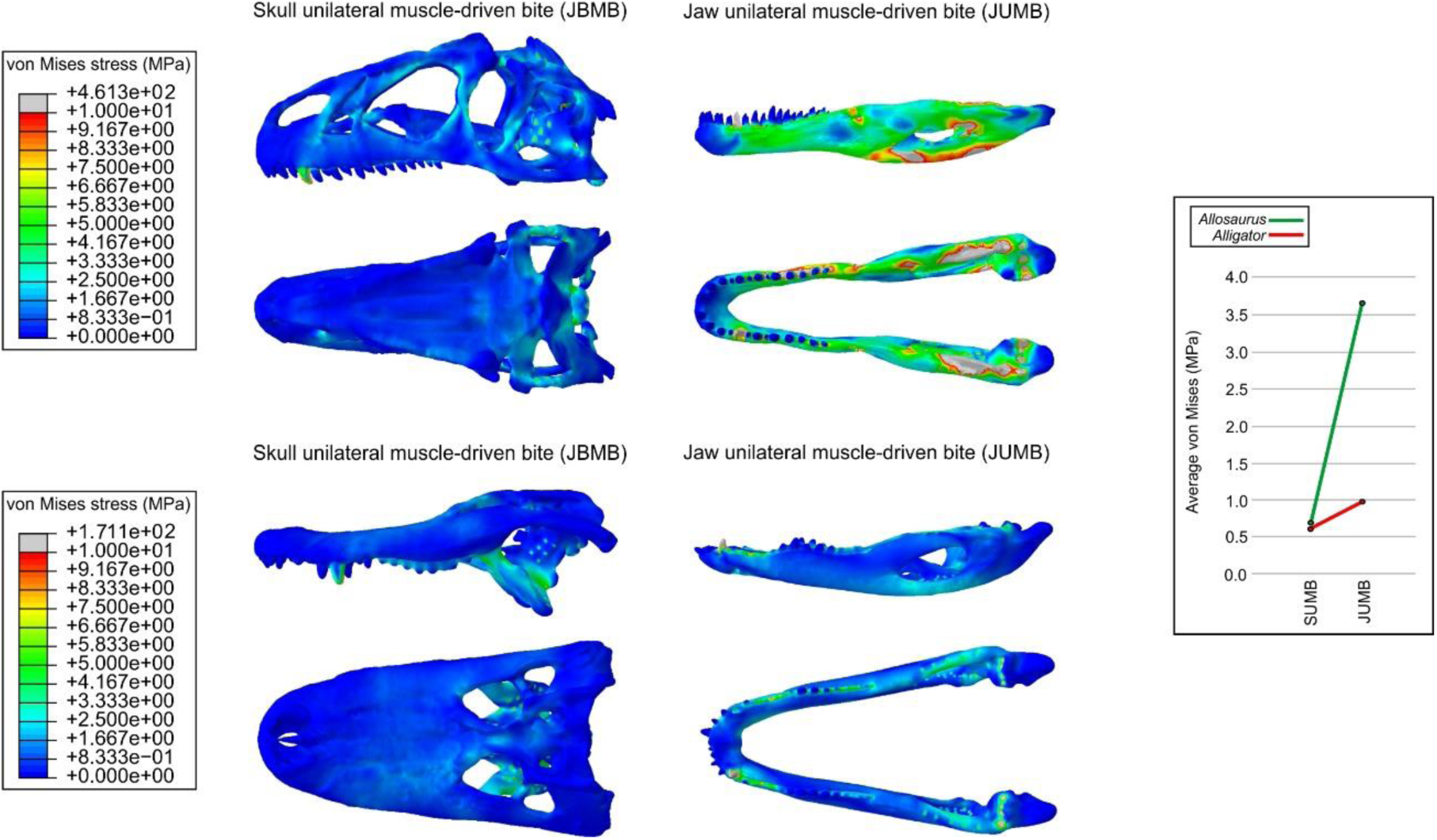
Von Mises stress contour plots from FEA of *Allosaurus fragilis* and *Alligator mississippiensis* for the intrinsic scenarios. Average Von Mises values per scenario for each taxon are displayed on the right.

**Figure 4.**
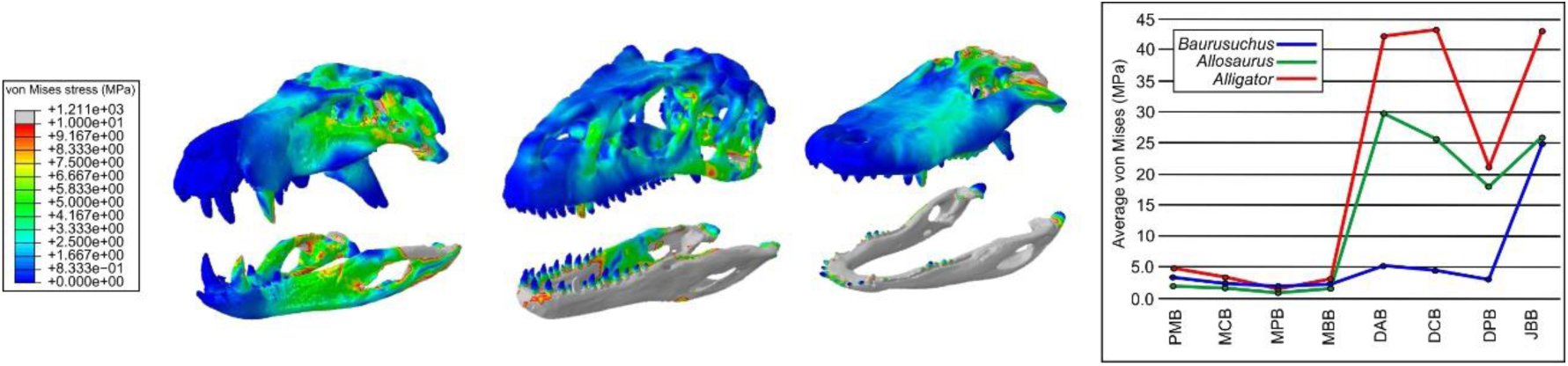
Comparison of Von Mises stress distribution for scaled models of different archosaurian carnivores: baurusuchid, *Allosaurus fragilis* and *Alligator mississippiensis*. Stress contour plots displayed for the anterior bending scenario. On the right, comparative average Von Mises values per scenario for each taxon.

The two taxa retrieved greater differences in the lower jaw models during the intrinsic scenarios (average Von Mises stress of 3.7 MPa for *Allosaurus* and 0.99 MPa for *Alligator*). The discrepancies observed in the bending scenarios are also most evident in the lower jaws, which for the baurusuchid remain consistently less stressed than those of both the theropod and the crocodylian during the bending tests. When compared to the baurusuchid, the theropod models obtained only slightly lower average Von Mises stress values for the skull, but much higher values for the lower jaws (Figure 4). The *Alligator* model, in contrast, retrieved higher average Von Mises stress values in most scenarios than both the baurusuchid and *Allosaurus*, even though differences in stress values are less distinguishable between skull models of the analysed taxa (Figure 4). The only scenario that does not follow this pattern is the unilateral bending at the back of the upper-tooth row, in which the average Von Mises stress value is similar for the baurusuchid and *Alligator*, although both have higher stresses than the theropod. The most divergent results are related to the mandibular anterior bending scenario, in which the average stress value in *Alligator* was more than nine times higher than in the baurusuchid, and almost twice the average Von Mises stress recorded for the theropod.

## DISCUSSION

The unexpectedly weak bite force estimated for the baurusuchid is much lower than that measured for extant crocodylians of comparable size (*Alligator sinensis* with a total body length around 150 cm can have a bite of up to 963N measured at the caniniform tooth, Erickson *et al*. 2012). It is also only a fraction of the bite forces inferred for adult theropods, which could potentially exceed 50,000 N (Gignac & Erickson 2017). This comparatively weak bite force in baurusuchids suggests that their role as apex predators may have involved hunting strategies different from those of most carnivorous theropods and living crocodylians, which mostly rely on muscle-driven biting forces for killing (Rayfield 2004, 2005, 2011; D’Amore *et al*. 2011; Erickson *et al*. 2012). As a consequence, the killing potential of baurusuchids could have been enhanced by structural and behavioural traits, as in other weak-bite apex predators such as troodontids and allosaurid theropods, varanid lizards and felines, that harness the post-cranial musculature to supplement bite force (Rayfield 2001; D’Amore *et al*. 2011; Figueirido *et al*. 2018; Torices *et al*. 2018).

Alternatively, the apex predator role of baurusuchids could have been a historical misinterpretation, and the group might be better interpreted as preying on smaller and/or softer animals. However, a series of craniomandibular and postcranial adaptations of baurusuchids indicate otherwise. For example, the presence of extensive overengineered regions around the canines in both the skull and lower jaws (e.g. regions that remain relatively stress-free in all tests) show that the baurusuchid craniomandibular architecture could safely perform in much higher stress conditions than imposed by muscle-driving biting forces. This is true even for our bending tests that most likely overestimate the stress experienced by the skull of the baurusuchid. The presence of overengineered regions in *Allosaurus* has been suggested as evidence that this taxon also used mechanisms to enhance killing potential in its regular feeding strategy (Rayfield *et al*. 2001).

Additionally, the tested pull-back, head-shake and head-twist scenarios were designed to understand how the baurusuchid craniomandibular architecture would perform during similar head movements employed by other weak- and strong-bite apex predators (Rayfield 2001; D’Amore *et al*. 2011; Torices *et al*. 2018). For baurusuchids, these movements would be possible given the robust cervical vertebrae, high neural spines, and well-developed cervical ribs (particularly the first two), which provided large attachment areas for the muscles responsible for head lift, head twist, and side-to-side movements (Cleuren & De Vree 2000; Godoy *et al*. 2018). These tests show that the baurusuchid skull and mandible worked optimally in scenarios simulating non-orthal loads, suggesting that baurusuchids were well-suited for head movements during predation, possibly even more than living crocodylians. This can be explained by the combination of three skull features that minimize skull stress during bites and torsion, the oreinirostral morphology, the absence of the antorbital fenestra, and the extensively ossified secondary palate. This combination of features is particularly efficient for stress reduction during unilateral biting (Rayfield & Milner 2008).

Our tests also revealed that the well-developed gap between premaxillae and maxillae is a unique specialization in the skull architecture of baurusuchids, very likely related to predatory habits. This gap redirects the stress from the premaxillae to the dorsal surface of the fused nasals during biting, preventing stress from traveling from the occlusal region of one bone to the other, and implying a functional decoupling between premaxillae and maxillae during bites. This gap at the premaxillae-maxillae suture is absent in *Allosaurus* and *Alligator*, and in those taxa, the stress travels directly from the premaxilla to the maxilla, especially during the unilateral premaxillary bending scenarios. A similar stress redirection is observed in tyrannosaurids, in which the robust and also fused nasals work as main route for stress distribution, bypassing the less robust maxilla-lacrimal contact (Rayfield 2005). We suggest that the gap observed in baurusuchids, in combination with the robust and fused nasals, worked similarly to that of tyrannosaurids, even though, the general cranial architecture presented by the baurusuchid is closer to the *Alligator*. The gap could also allow repeated punctures to be inflicted from biting at different positions of the tooth row, but concomitantly working as a built-in safety factor, minimizing the risk of the skull yielding (Rayfield *et al*., 2001). Finally, the presence of ziphodont dentition in baurusuchids is also in line with the role of apex predator (Riff & Kellner 2011; Godoy *et al*. 2014). Knife-like teeth with well-developed serrated cutting edges are a dental adaptation for optimal defleshing of vertebrate carcasses (D’Amore *et al*. 2009) and are present in a series of unrelated apex predators, including theropod dinosaurs and large monitor lizards (D’Amore *et al*. 2011; Brink & Reisz 2014; Torices *et al*. 2018).

The discrepancy in the Von Mises stress magnitude and distribution seen between the mandibles of the three taxa during the intrinsic scenarios and during the bending tests suggests that this structure is also pivotal in understanding the palaeoecology of baurusuchids. The Von Mises stress distribution shows that *Allosaurus* and *Alligator* have, in general, higher and more homogeneously distributed Von Mises stress in the mandible, while in the baurusuchid the stress is concentrated at the postsymphyseal region. This indicates that the robust symphysis in baurusuchids is important for stabilizing the lower jaws.

The best example of the divergent responses among lower jaws is seen in the bilateral bending scenario, for which the average Von Mises stress value for the baurusuchid was approximately five times greater than any other scenario. Additionally, this is the only scenario in which the Von Mises stress approaches the higher values presented by *Allosaurus* and *Alligator* (Figure 4). The baurusuchid response is also different from *Allosaurus* and *Alligator* in the sense that the average Von Mises stress values in the bilateral bending scenarios are distinct from the unilateral scenarios, whereas the other two taxa show similar values in both scenarios. Based on our FEA results, we propose that the bilateral biting is the least likely killing strategy for baurusuchids, and the clamp-and-hold, employed by living crocodylians, and large mammal predators, such as the lion (*Panthera leo*) (Figueirido *et al*. 2018), does not fit the mechanical properties of the baurusuchid skull.

Our results also indicate that baurusuchids were well adapted for handling struggling prey, which was possibly subdued by inflicting a series of bites using premaxillary, maxillary and particularly the dentary canines, that combined with ziphodonty would pierce repeatedly the skin of the prey. The puncture phase would be followed by head-movements that would worsen the wounds caused by the punctures and ultimately leading to the death of the prey.

Our results successfully characterise the exceptional suite of biomechanical properties displayed by baurusuchids, which combine novel adaptations, features similar to theropods, and others seen in living crocodylians. Such a combination has not been reported previously for any predatory taxon, raising questions on the specific evolutionary settings that allowed these features to emerge. Selective pressures from extrinsic environmental factors seem to have an important influence during amniote functional and biomechanical evolution (Sakamoto *et al*. 2019). In the case of baurusuchids, the unique Late Cretaceous palaeoecosystems of southeast Brazil exhibited a combination of playa-lake systems and transitory rivers which possibly permitted life to flourish in semi-arid to arid conditions (Carvalho *et al*. 2010; Marsola *et al*. 2016). These landmasses witnessed an extraordinary diversity of crocodyliforms (especially notosuchians; Mannion *et al*. 2015), as well as other tetrapods (Godoy *et al*. 2014). This resulted in a diverse array of potential prey for baurusuchids among terrestrial crocodyliforms, indicating that prey selection could have played an important role in the evolution of the baurusuchid craniomandibular apparatus.

## ACKNOWLEDGEMENTS

This work was supported in part by a Rutherford Fund Strategic Partner Grant to the University of Birmingham, which funded the travel of XXXX to Birmingham. This research was supported by a National Science Foundation grant (NSF DEB 1754596) to XXXX and Fundação de Amparo à Pesquisa do Estado de São Paulo (FAPESP 2019/10620-2) to XXXX.

